# Analysis of the Influence of Hypergravity State in the Simulated Aerospace Flight Environment on the Cardiac Structure of Amphibians during Development

**DOI:** 10.1101/2023.04.18.537410

**Authors:** Wang Jiahao, Li Zheyuan, Wang Yidi, Liu Cenxiao, Hu Aihua, Rehman Haseeb Ur, Wang Danmei, Wang Yang

## Abstract

**Background:** Hypergravity environment is a kind of extreme environment that human beings will inevitably encounter when they realize space navigation. When the body is affected by a hypergravity load, the instantaneous changes in fluid distribution cause abnormalities in the physiological functions of the heart and blood vessels. Whether to adapt to these extreme conditions is an important link for humans to break through the Earth’s exploration of space.

**Method:** This study adopts the experimental method of simulating hypergravity, using amphibian frog larvae as the research object, to observe the structural changes of the unique single ventricle of amphibian frog larvae after being subjected to hypergravity load. Combining digital simulation technology, this study explores the possible impact of hypergravity load on ventricular function. The experiment selected frog larvae (*Larvae*, commonly known as tadpoles) and subjected them to a continuous load of 10 minutes under a rotating supergravity state of +3Gz for 3wks. The hypergravity load experiment ends when the larvae develop into young frogs (*Metamorphs*). After the specimen is subjected to histochemical fixation treatment, it is then embedded, sliced, stained, and subjected to computer-assisted microscopy to obtain heart slice images. With the help of computer-assisted image analysis, the length, axis, and ratio of the ventricles are calculated, and the morphological changes of the ventricles are analyzed.

**Results:** Research shows that the impact of hypergravity fields on the heart is multifaceted. Due to prolonged and intermittent hypergravity load stimulation, the swimming mode of juvenile frogs has changed from a normal symmetrical swing of the tail to a dominant swimming mode on one side. The vestibular nucleus discharge record shows that after hypergravity load, the activity of vestibular nucleus discharge in juvenile frogs is lower than that in the control group, indicating that simulated hypergravity load has an effective stimulating effect on the development of amphibian frogs from larvae to juveniles. Hypergravity also causes the heart to shift to the right within the chest cavity, resulting in elongated ventricles with an imbalance in the ratio between the longitudinal and transverse axes, indicating a possible decrease in filling capacity.

**Conclusion:** The experimental results of this study suggest that the hypergravity loading environment during space navigation can affect ventricular structure, and changes in this structure can reduce cardiac ejection function. Starting from the conclusion that prolonged intermittent hypergravity loads can affect heart development, it is necessary to consider how to develop protective equipment to alleviate the thoracic space bearing hypergravity loads, reduce cardiac anatomical displacement and ventricular structural imbalance, and ensure that the body maintains normal cardiac blood supply function in the airspace environment. This is a topic that needs further exploration in the future.

## Introduction

2025 is the first year for humans to embark on manned space flight. China will send the “Zhenghe” spacecraft to land on an asteroid to collect samples that year, while the US National Space Administration has repeatedly legislated in 2010 to achieve the goal of sending humans to an asteroid by 2025 and to *Mars* by 2030. In order to achieve this dream of humanity, the research and development of manned spacecraft is in the ascendant period of development and growth. The space medicine associated with it is becoming increasingly important as a necessary condition for future safety and successful mission execution.

In the academic seminars on space environmental medicine attended by Chinese and foreign key medical students in the international undergraduate program of clinical medicine in our hospital in December 2022 and March 2023, we have realized that cardiovascular physiological function is a subject that needs to be discussed in depth in human space missions. In the process of space flight, human beings need to overcome the two very special gravity field environments of hypergravity and microgravity, This is also the challenge of cardiovascular physiology in the extreme environment faced by the current manned space program. Early scientific research using mammals as research subjects found that mammals can tolerate and adapt to sustained hypergravity of up to 2.5g without seriously damaging their body structure and function ^[1]^. Research on human body shows that when hypergravity increases to three times of the normal value, human can significantly increase cardiac output through lower limb movement, and the increase rate is significantly higher than the increase rate of heart rate ^[2]^.

The method of underwater immersion or posture change is usually used to simulate the hypergravity and microgravity of space environment. The simulation experiment shows that when the body is exposed to 0g (water immersion or a head down tilt position of - 6 degrees) and the head to foot positive acceleration is implemented when the high gravity is eliminated and the micro gravity is transferred, there is a specific functional correlation between body fluid and cardiac output ^[3]^. When in a low body negative pressure (LBNP) state with head low and legs high, it is equivalent to being in a hypergravity state of +2Gz, and there will be a significant change in heart rate (HR). Both upright and supine states of the body in LBNP state can have an impact on hemodynamic indicators. During upright state, HR changes exceed the HR level during supine state, leading to an increase in heart rate. However, HR peaks when supergravity reaches +3Gz and +4Gz, indicating that a change in posture can interfere with the study of cardiovascular function in a hypergravity environment, thereby affecting the adaptive response of cardiovascular function, the regulatory mechanism of the circulatory system, and interfering with hemodynamic parameters, Amphibians, on the other hand, do not suffer from upright or recumbent postures during their larval stage, making them highly suitable for studying cardiovascular adaptability in high-gravity environments.

The changes in the distribution of hydrostatic pressure in the body caused by hypergravity can cause changes in the arterial system and venous reflux of the circulatory system. In the physiological state of normal body, the left cardiac output is about 2% higher than the right heart. Although there is no significant difference in statistics, this small difference has important physiological significance in maintaining the dynamic balance of blood flow between the left heart’s systemic circulation and the right heart’s pulmonary circulation. When the instantaneous gravity changes, there is asymmetry between the left ventricular cardiac output (QLV) and the right ventricular cardiac output (QRV) ^[4]^, the blood flow balance between the left and right hearts is broken, and the mechanical load borne by the myocardium is out of balance, which may have an impact on the cardiac structure during development. On the other hand, the blood volume in the pulmonary vessels showed an instantaneous increase, and the level and direction of hypergravity had a great impact on the increase amplitude and duration. Especially in the period after the end of hypergravity and hypergravity, the continuous instantaneous changes in the blood volume of the pulmonary vascular system put forward high requirements for the compliance of the pulmonary circulation system, and these changes increased the impact on the development of the right heart.

Research on the cellular level of myocardial cells has found that when exposed to a hypergravity state of +1.8G, both cardiac mass and mitochondrial volume show an increasing trend ^[5]^. At this time, hemodynamic indicators show a significant increase in systolic and diastolic blood pressure, and a narrower range of pulse pressure values ^[6]^. In terms of energy supply to the myocardium, hypergravity can cause a significant increase in Ca^2+^ transient in the myocardium. At this time, the energy consumption of the myocardium is higher, and the ATP consumption of the myocardium sharply increases, manifested by an increase in the content of inorganic phosphate (Pi) in the myocardium and a decrease in the concentration of phosphocreatine (PC).

Morphologically, it is characterized by a larger heart mass and mitochondrial volume. At this point, the hemodynamic indicators show a lower left ventricular end diastolic pressure (LVDP), and the myocardium exhibits a high compliance state, with an increase in mechanical length load of the myocardium. The significant changes in LVDP and Pi/PC ratio values indicate that the effect of hypergravity on the heart is to generate more energy expenditure ^[7]^. Cardiac output decreased by 20% - 50%, and cardiac function curve of CO and AP shifted to the right ^[8]^. The state of hypergravity can also affect the expression of gene levels in myocardial cells, leading to a significant decrease in the linear staining of the basement membrane area of developing myocardial tissue and a significant decrease in the number of desmosomes per unit area. This change greatly reduces myocardial contractile function ^[9]^.

The parabolic flight experiment is another large-scale high-gravity simulation experimental method. In the analysis of experimental results on 8-10 parabolic flight cycles, it was found that during continuous parabolic flight, factors such as pleural effusion index and cardiac index showed significant changes when the initially generated supergravity parabolic flight reached 6, 7, and 8 cycles ^[10]^. Significant changes were found in both right atrial pressure and left atrial pressure ^[11]^. During hypergravity phase I, the inflow velocity of the right ventricle during the filling phase increased by 46%, the left ventricular ejection velocity increased by 12%, and the inflow velocity during the filling phase increased by 13%. The experiment showed that the right ventricle was subjected to additional venous reflux, resulting in a significant increase in the mechanical load generated by ventricular blood volume ^[12]^. At a hypergravity state of+1.8Gz, the end diastolic (ED) and end systolic (ES) filling degree of the ventricle significantly decreases, the filling fraction increases, and the left ventricular filling caused by atrial contraction decreases ^[13]^. Long term exposure to an instantaneous hypergravity environment can result in mitral valve prolapse (MVP) but no significant reflux in the body ^[14]^. The changes in these hemodynamic indicators all point to the point where the heart undergoes significant mechanical loads during and after a period of hypergravity, which has a significant impact on the structure and function of the heart. The significant decrease in average arterial pressure caused by hypergravity will have a significant impact on cerebral blood flow and affect the regulation of the nervous system on the body ^[15]^. The study of the impact of heart development on the evolution of amphibian larvae into frogs during their developmental stages can reveal various abnormal changes in heart function that may occur under high gravity conditions, providing some inspiration for addressing cardiovascular maladaptation during space travel.

In terms of cardiac electrophysiology, which affects the automatic rhythm of the heart, the state of high gravity will lead to an increase in the occurrence of sudden cardiac arrest (SCA). Because of the cardiopulmonary resuscitation method adopted by manned spacecraft in space environment, the depth of the compression operation and the ability to continue external chest compression are lower than the earth’s standards, The changes in the position and structure of the heart caused by hypergravity also pose more special requirements for the automatic chest compression device (ACCD) [16].

By artificially simulating the hypergravity experimental environment, it can be used to discover and master clues for preventing or treating various cardiovascular physiological disorders during space flight. It can be further used for the research, development, and application of protective equipment.

Amphibians, especially frogs, have significant similarities with mammals, and the research conclusions obtained through frog experiments have been widely recognized by the international academic community. Experimental validation results have also been adopted by the Nobel Prize in Medicine evaluation system. Due to its ease of maintenance, it has been widely used in life science research in the aerospace field.

The launch of Bigelow Aerospace’s Genesis-1 in 2006 marked the world’s first inflatable space station, leaving ample space for physiological research on amphibians and other low gravity animals. Since 2020, the European Space Agency (ESA) has launched the Animal Model Gravity Experiment Platform (GEPAM) in its land-based facility portfolio to study the impact of gravity changes on amphibians. With the development of China’s spacecraft industry, animal experiments in outer space are also constantly being explored.

The development of amphibian frog larvae into adults can be divided into 10 stages (10 stages, **Figure 1**A). This study selected 5 stages of frog larvae (*Larvae*) as the experimental subjects. Amphibian juvenile frogs have a strong ability to regenerate after injury due to their unique single ventricular structure (**Figure 1**B) in the heart. The frog’s heart has a special physiological regulatory ability to regulate pumping function by changing the ventilation mode of the respiratory cycle, which has attracted attention from cardiovascular medical researchers. Its myocardium and mammals have the characteristics of easy observation and good experimental repeatability in terms of myocardial cell excitability (especially the mechanism of Ca^2+^ transient) and contractile force function. This experiment focuses on observing the impact of hypergravity load on the development and growth of the ventricles of larvae from stage 5 to stage 8, and evaluating the structural characteristics of the ventricles that are subjected to hypergravity load and produce compliance.

**Figure 1.**
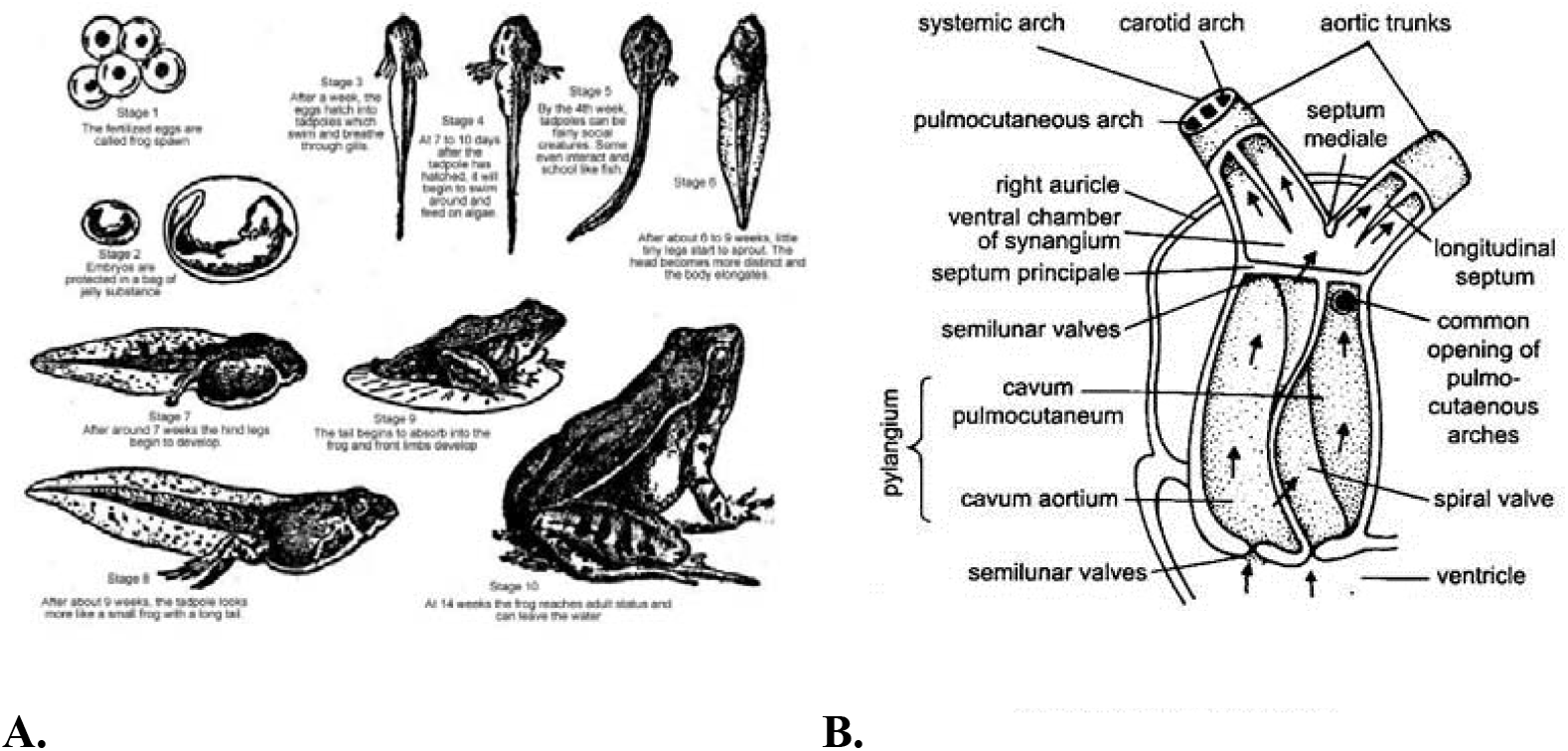
The body shape changes in hypergravity simulation juvenile frogs.

## Materials and methods

### Experimental subjects

This experiment was conducted in accordance with the notice of the Provincial Department of Science and Technology on strengthening the biosafety management of experimental animals in our province, in accordance with the “Regulations on the Management of Experimental Animals” and in combination with our college’s regulations on the management of experimental animals, under the guidance of technical teachers in the functional laboratory. During the experiment, the operating procedures for experimental animals in the functional laboratory were strictly followed.

The experimental animals selected were amphibian frog *larvae* (5 stages), which were raised in a specially designed water tank in the laboratory. Their growth was controlled by day and night light at room temperature for 12 hours/12 hours, and each individual was fed in separate compartments for 1week (1wk). After normal feeding, a 3wks hypergravity simulation experiment was conducted. All experimental subjects underwent static planktonic observation twice a day in the morning and evening to ensure their viability.

### Equipments

The rotating sleeve, specially designed water tank, and related auxiliary equipment used in the experiment of simulating hypergravity state with horizontal angular velocity movement are all provided by the Functional Laboratory of the Medical College. The experiment used a manual drum to measure the diameter of the rotating drum before use, and calculated the correlation between the rotating speed of the drum and the hypergravity generated by the outer diameter edge of the drum. The final speed was set to 30 times/min at 3GHz.

The numerical simulation software and image acquisition equipment for cardiac hemodynamics are provided by the functional laboratory.

The video imaging equipment in the static planktonic experiment is provided by the functional laboratory; The specimen preparation technology, slicing, staining, and morphological imaging equipment for the later stage of the experiment are provided by the Morphology Laboratory of the Medical College.

### Hypergravity Simulation

#### 1. Horizontal rotational motion

The angular velocity movement experiment is a 3wks experiment conducted in our institute’s functional experiments. The rotating sleeve and auxiliary equipment are provided by the functional laboratory. The amphibian frog larvae (n=20) were randomly divided into a hypergravity load group (n=10) and a control group (n=10) after 1wk of normal feeding. The larvae of the hypergravity load group were placed in a water tank set in the rotating drum and subjected to a hypergravity load of+3Gz at a speed of 30 times/min. After 10 minutes, they were transferred to the feeding tank twice a day in the morning and evening. This operation was repeated daily for a total period of 3wks.

After implementing a load of 2wks, half of the larvae were harvested, and the remaining portion continued until 3wks. The control group larvae were fed normally (n=10) without applying hypergravity load.

Before and after each hypergravity load, video recording of larval activity and swimming activity, while observing their body length and limb growth.

#### 2. Planktonic experiment

Place the larvae of the hypergravity load group and the normal group in a horizontal water tank, record the swimming situation of each group in the water through video, analyze and compare the floating and swimming motion curves caused by changes in the neural motor regulation system of each group through video imaging.

### Specimens Collection

After the hypergravity loading experiment, the young frogs (Metamorphs) were fixed in a 4% formaldehyde solution and embedded in paraffin. Perform 7 μ m slicing on the horizontal plane of the young frog. H. E. After staining, the horizontal anatomical images of the chest of the young frog were taken under an optical microscope.

### Myocardial tissue image acquisition and computer-aided analysis

NIH ImageJ software (Java 1.8.0_122) use for perform semi quantitative analysis of ventricular images captured under an optical microscope.

### Analysis of ventricular structure data

Due to the unique anatomical structure of a single ventricle in amphibian juvenile frogs, changes in fractional area have been shown to have a good correlation with the ventricular function index derived from the pressure volume loop. This experiment calculates the long and short axis of the ventricles, as well as the ratio between them, based on the ventricular images obtained from the slices of young frog specimens, and compares the statistical differences between the hypergravity load and the control group.

### Statistical Analysis

The experimental data results were mean ± standard error (Mean ± SEM), and Mann Whitney U test was used to compare the differences in ventricular morphology between the+3Gz hypergravity load group and the control group after two months. P<0.01 indicates a statistically significant difference.

## Results

After a total period of 3wks of implementation of the hypergravity load, both the hypergravity group and the control group larvae developed into juvenile frogs. Its characteristic is normal limb development, with significant increase in weight and length. However, some larvae in both groups died prematurely due to unadaptation to artificial feeding environments.

After implementing a loading cycle of 2wks, the survival rate of the hypergravity composition was 80% (8 survived). Three *larvae* were taken for specimen preparation, and the remaining 5 *larvae* were maintained in the hypergravity loading experiment until 3wks; The survival rate of the control group for 2wks was 90% (9 survived), and 3 were taken for specimen preparation, while the remaining 6 were kept in normal feeding until 3wks. After 3wks, there were 4 live animals in the hypergravity group (n=4) and 5 live animals in the control group (n=5) (**Table 1**).

**Table 1.** The body length and weight after 3wks hypergravity simulation.

| Grouping | Before |  |  | After ( 3wk ) |  |  |
| --- | --- | --- | --- | --- | --- | --- |
|  | Body length | Body weight | BMI | Body length | Body weight | BMI |
| Simulation Group | n=10 |  |  | n=4 |  |  |
|  | 5.10±0.18 | 27.80±0.93 | 1.16±0.10 | 35.80±0.81 | 7.20±0.12 | 0.72±0.04 |
| Control Group | n=10 |  |  | n=5 |  |  |
|  | 4.70±0.26 | 27.00±0.89 | 1.30±0.12 | 36.76±0.94 | 8.30±0.45 | 0.78±0.05 |

After 3wks of loading, the juvenile frog in the hypergravity load group showed a right-sided bending posture in the tail during resting state, while the control group showed normal behavior.

### Posture changes in simulated juvenile frogs

In the hypergravity load group, intermittent hypergravity load resulted in a one-sided dominant movement pattern in the developing young frogs. As shown in **Figure 2**, the planktonic body state of the *larvae* before the experiment in A, B, C, and D are swimming bodies of 2wks and 3wks, respectively. The main manifestation is that the head deflects to the left when in a quiet state, while the tail bends to the right. The swing amplitude of the tail to the right during swimming is significantly greater than that to the left, and the extensor muscle of the left limb is more active than that of the right limb during swimming.

**Figure 2.**
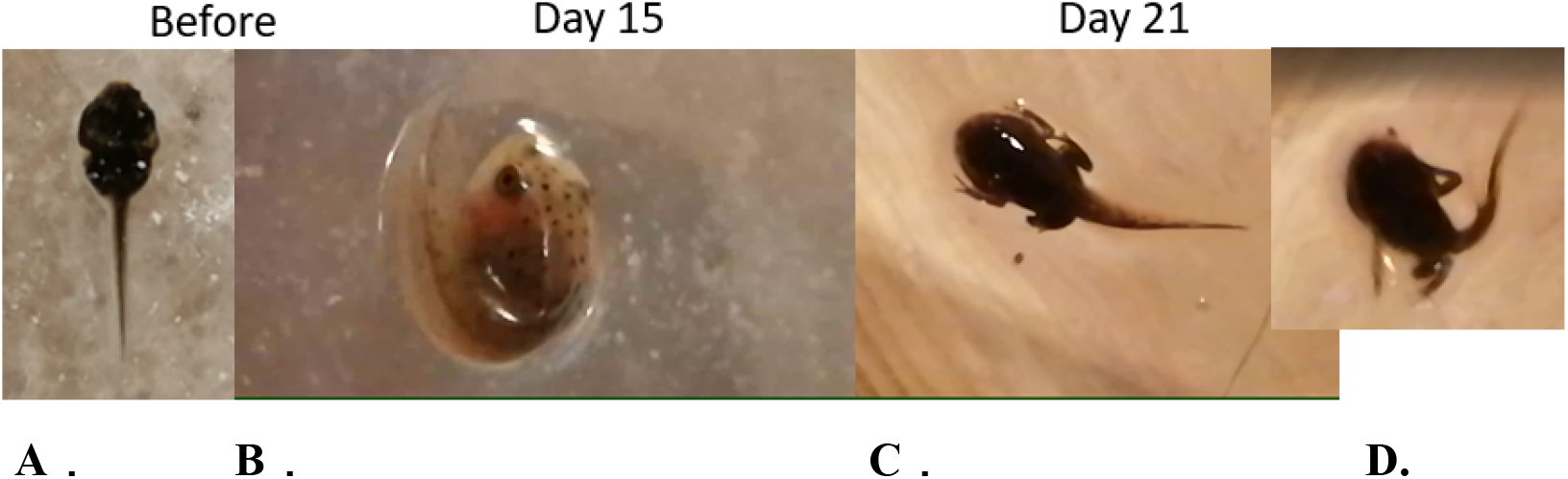
The body posture changes in simulation juvenile frogs (dorsal view)

Observing the young frog from the back, it was found that its paddling movement exhibited a left-handed advantage in the opposite direction of rotation to the direction of rotation when applying hypergravity load (clockwise rotation when applying hypergravity load).

### Vestibular nucleus discharge changes after simulation

The intermittent hypergravity load group of young frogs caused a decrease in the activity of vestibular nucleus discharge and delayed waveform of vestibular nucleus discharge induced by stimulation of lower limbs (**Figure 3**).

**Figure 3.**
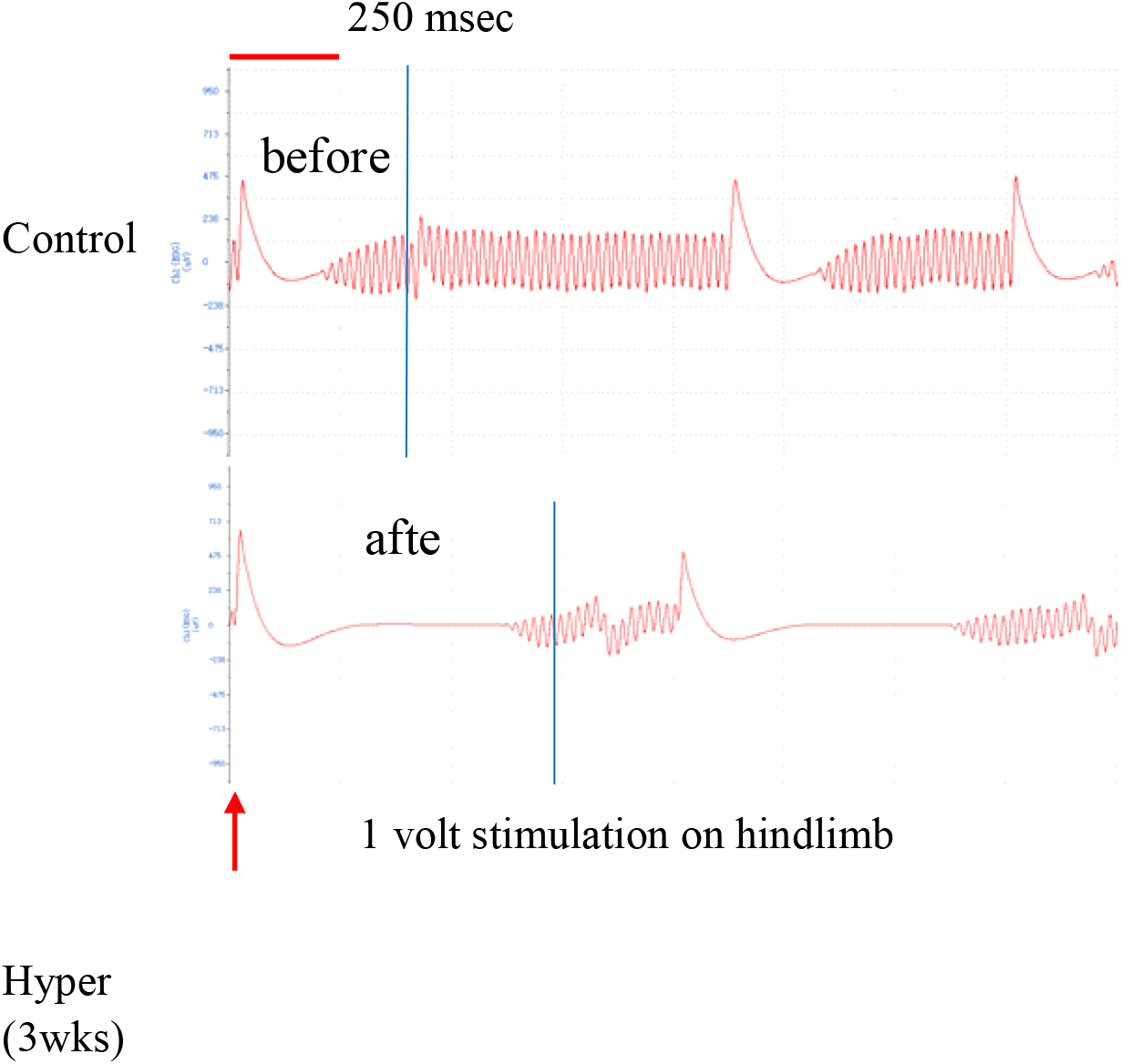
The delayed discharge of the vestibular nucleus in juvenile frogs induced by the lower limbs of the frog after 3wks simulation.

### Cardiac anatomic postion changes after simulation

After intermittent hypergravity loading, the dorsal horizontal section of the young frog showed that its overall heart position shifted to the right, and it was more pronounced after a total cycle of 3wks of loading. The heart structure of young frogs after each cycle of hypergravity loading is intact, specifically manifested as the complete ventricular wall structure of larvae after 2wks of hypergravity loading (**Figure 4**A); After 3wks period of hypergravity loading, the heart of the young frog still has a complete structure of two atria and one ventricle, with clear pericardial structure and obvious atrioventricular sulcus structure (as shown by the arrow in **Figure 4**B).

**Figure 4.**
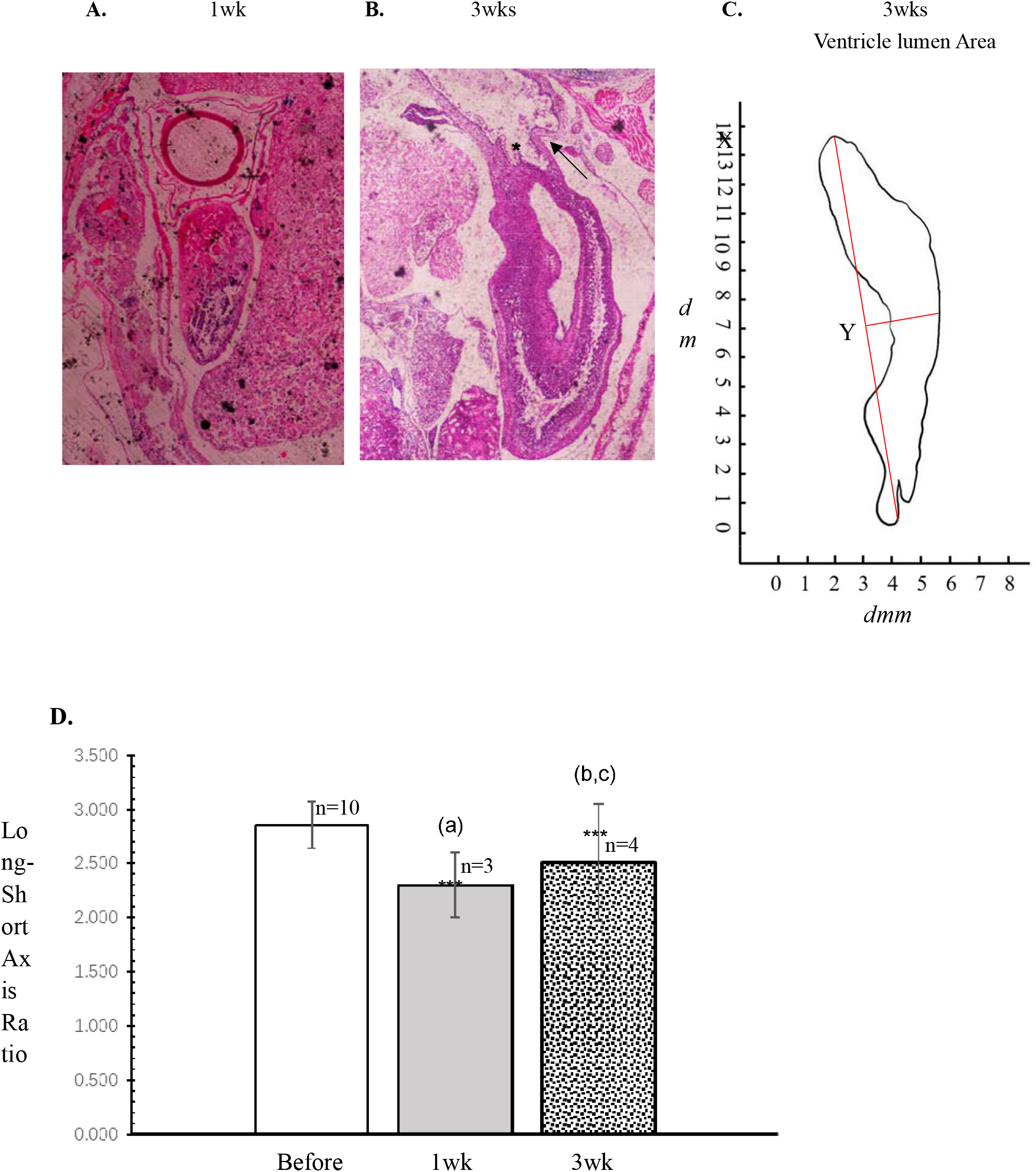
Cardiac ventricular structure after simulation.

Due to the hypergravity load caused by the clockwise angular velocity movement, the anatomical position of the heart of the young frog after a total cycle of 3wks leans to one side within the chest cavity, and the entire ventricle presents a slender state, with the ventricular wall bending to the right.

### Cardiac ventricular structure changes after simulation

After implementing a total cycle of 3wks of loading, the ventricle remains a single sponge like ventricular cavity, with a complete ejection filling structure at a single outlet (**Figure 4**B, *), and a movable spiral structure similar to a valve. The complete cavernous ventricle and valvular spiral structure in the arterial cone indicate that the young frog still has the physiological structure of balancing systemic circulation and pulmonary circulation after long-term hypergravity loading.

There were significant statistical differences in the aspect ratio of the single ventricular chamber between the *larvae* with a period of 2wks of hypergravity loading and the larvae with a total period of 3wkss after loading compared to the control group before loading. This result suggests that as the accumulation time of hypergravity load increases, the ventricular structure shows significant changes. When comparing the longitudinal and transverse axis ratios of the ventricular cavity between 2wk and 3wk, it was found that the longitudinal and transverse axis ratios of 3wk were significantly higher than those of 2wk larvae, indicating a qualitative change in the development of the ventricle after a continuous 2wk. **Figure** 4D shows a comparison between the hypergravity group (n=3) and the control group (n=3) after 2wkss of implementation; B and c are the hypergravity group (n=4) and control group (n=5) after 3wk, as well as the 3wk hypergravity group (n=4) and 2wk hypergravity group (n=3), respectively. * * * is p<0.01.

## Discussion

This study shows that the impact of hypergravity environment on the heart is multifaceted. Under long-term intermittent hypergravity load environment, the swimming mode of young frogs has changed from normal symmetrical swinging of the tail to a dominant swimming mode on one side. The electroencephalogram showed that after the implementation of hypergravity load, the discharge activity of the vestibular nucleus in the juvenile frog was lower than that in the control group, indicating that the simulated hypergravity load had an effective stimulating effect on the development of larvae into the 5th to 8th stages of the juvenile frog.

This study also found that long-term intermittent hypergravity loading can affect the normal development of the viscera, which first manifests as the normal anatomical position of the viscera shifting to one side. Under the overload environment caused by long-term clockwise rotation acceleration, the heart position significantly shifts to the right.

In addition, the study also found that there were significant morphological changes in the ventricles of young frogs after implementing a total load cycle of 3wks, mainly resulting in 1) elongated changes in the ventricles and a semi lunar curvature, with significant changes in the aspect ratio of the ventricular cavity. This may be related to the redistribution of fluid within the thoracic cavity caused by the clockwise needle angle variable hypergravity load and the impact of pressure generated by visceral interactions on cardiac development. This structural change in the ventricle is most pronounced after 3wks of load implementation.

Computer assisted image processing shows that there is a significant increase in both the longitudinal and transverse axes of the ventricular cavity after 2wks of intermittent hypergravity loading, while the ratio of the longitudinal and transverse axes decreases significantly. This indicates that the development of the ventricle in the transverse axis can still maintain a certain level during the 2wks of intermittent hypergravity loading. However, during the implementation period of intermittent hypergravity load for 3wks, the development of the ventricles in the longitudinal axis direction was significantly higher than that in the transverse axis direction after this prolonged intermittent load, and the ratio of the longitudinal and transverse axes of the ventricular cavity showed a very significant increase. The experimental results suggest that there is a positive correlation between the impact of hypergravity load on the development of ventricular structure and the length of the loading cycle. Even if short-term hypergravity load is intermittently applied (10 minutes), if this stimulation is repeatedly applied for a long time, it will have a significant impact on the variation of ventricular structure.

The aspect ratio of the ventricles affects ventricular blood filling and ejection. Especially the long axis of the ventricular cavity, it is an important indicator for determining ventricular filling and emptying index in clinical imaging medicine. The long axis of the ventricular cavity is also an important diagnostic indicator for determining the mechanical contraction mechanics of ventricular muscles, and has important reference value for the development of natural myocardial disease and clinical changes in cardiac structure and function ^[17]^. The difference between the maximum and minimum lengths of the long axis during the same cardiac cycle reflects the level of ventricular systolic function ^[18]^. Prolonged long axis of the ventricular cavity can lead to a decrease in the difference between the maximum and minimum lengths, indicating a decrease in ventricular diastolic function, limited blood filling, and a decrease in ventricular stroke volume. The results of this experiment suggest that in the early stages of space navigation, the short-term hypergravity load generated by overcoming Earth’s gravity mainly affects the heart after periodic stimulation, with an increase in the long axis of the ventricle, a decrease in ventricular filling and ejection capacity. It is necessary to further develop new protective devices that can protect against periodic hypergravity stimuli in the future, in order to address the negative effects on heart function caused by repeated shuttling between Earth and space. This study provides new clues for the further development of new space suits and the absorption of new technologies to protect the chest from the effects of hypergravity loads, and provides valuable basic medical research clues for solving the adverse factors of long-term space shuttle on the body.

